# Spectral neural approximations for models of transcriptional dynamics

**DOI:** 10.1101/2022.06.16.496448

**Authors:** Gennady Gorin, Maria Carilli, Tara Chari, Lior Pachter

**Affiliations:** Division of Chemistry and Chemical Engineering, California Institute of Technology, Pasadena, CA, 91125; Division of Biology and Biological Engineering, California Institute of Technology, Pasadena, CA, 91125; Department of Computing and Mathematical Sciences, California Institute of Technology, Pasadena, CA, 91125

## Abstract

The advent of high-throughput transcriptomics provides an opportunity to advance mechanistic understanding of transcriptional processes and their connections to cellular function at an unprecedented, genome-wide scale. These transcriptional systems, which involve discrete, stochastic events, are naturally modeled using Chemical Master Equations (CMEs), which can be solved for probability distributions to fit biophysical rates that govern system dynamics. While CME models have been used as standards in fluorescence transcriptomics for decades to analyze single species RNA distributions, there are often no closed-form solutions to CMEs that model multiple species, such as nascent and mature RNA transcript counts. This has prevented the application of standard likelihood-based statistical methods for analyzing high-throughput, multi-species transcriptomic datasets using biophysical models. Inspired by recent work in machine learning to learn solutions to complex dynamical systems, we leverage neural networks and statistical understanding of system distributions to produce accurate approximations to a steady-state bivariate distribution for a model of the RNA life-cycle that includes nascent and mature molecules. The steady-state distribution to this simple model has no closed-form solution and requires intensive numerical solving techniques: our approach reduces likelihood evaluation time by several orders of magnitude. We demonstrate two approaches, where solutions are approximated by (1) learning the weights of kernel distributions with constrained parameters, or (2) learning both weights and scaling factors for parameters of kernel distributions. We show that our strategies, denoted by kernel weight regression (KWR) and parameter scaled kernel weight regression (psKWR), respectively, enable broad exploration of parameter space and can be used in existing likelihood frameworks to infer transcriptional burst sizes, RNA splicing rates, and mRNA degradation rates from experimental transcriptomic data.

**Statement of significance:** The life-cycles of RNA molecules are governed by a set of stochastic events that result in heterogeneous gene expression patterns in genetically identical cells, resulting in the vast diversity of cellular types, responses, and functions. While stochastic models have been used in the field of fluorescence transcriptomics to understand how cells exploit and regulate this inherent randomness, biophysical models have not been widely applied to high-throughput transcriptomic data, as solutions are often intractable and computationally impractical to scale. Our neural approximations of solutions to a two-species transcriptional system enable efficient inference of rates that drive the dynamics of gene expression, thus providing a scalable route to extracting mechanistic information from increasingly available multi-species single-cell transcriptomics data.

## 1 Introduction

The production, processing, and degradation of RNA molecules are governed by a set of stochastic events that lead to variable levels of RNA transcripts and proteins between genetically identical cells in identical conditions [1]. The inherent stochasticity and resulting variability in RNA transcript levels play an essential role in cellular function, development, and response, but must also be tightly regulated to produce deterministic functional outcomes at precise times. Extracting the rates that underlie distributions of RNA molecules is essential for understanding the intricate balance between stochastic gene expression and precise cellular responses required for life.

To explore the dynamics of transcriptional systems, specific biophysical models can be proposed that are explicitly parameterized by relevant biological rates, such as rates of transcriptional initiation, RNA splicing, or RNA degradation. These stochastic systems are naturally modeled by discrete-valued, continuous-time Markov chains (CTMCs). The evolution of a system’s microstate probabilities under a CTMC model of biology is given by a chemical master equation (CME) [2]. For example, a CME tracks the changes in probability of there being a certain number of nascent (*n*) and mature (*m*) RNA molecules at time *t* given certain biophysical parameters *θ*, or *P* (*n, m, t*; *θ*). This can be used to obtain insights into transcriptional biophysics [3] by fitting *θ* most consistent with experimental distributions of molecule numbers [4].

Until recently, mechanistic modeling and insight into the underpinnings of transcriptional processes have been tied to intensive, temporal, fluorescence datasets which generally measure the production of one species of RNA molecules in one cell line [ 1,5,6]. Recent high-throughput transcriptomics methods, such as single-cell RNA sequencing (scRNA-seq), offer discrete molecular readouts at genome-wide scale across diverse cell populations [7], thus making possible the exploration of transcriptional parameters across distinct cell-types, tissues, and organisms. In addition to increased cell-type coverage, the diversity of molecular species captured, such as the simultaneous quantification of unspliced and spliced RNA molecules [8–10], proteins [11], and chromatin accessibility [12], has the potential to deepen our understanding of transcriptional system dynamics [13]. While single-cell RNA-seq data has been modeled using CME solutions to extract biophysical rates of transcriptional initation and burst sizes, mechanistic modeling has not been adopted in recent analyses of multi-species transcriptomic experiments [14]. One major challenge is that closed-form solutions to CMEs that model multi-species systems with several interacting species are rarely available [3, 15], and current methods to solve them numerically for probabilities are prohibitive to the recursive calculations necessary for maximum likelihood inference of biophysical parameters.

The methods currently used to circumvent the lack of closed-form solutions to CMEs are computationally expensive and compromise accuracy. For example, CTMCs are often simulated using Gillespie’s stochastic simulation algorithm [16], which generates statistically realistic molecular trajectories over possible state-spaces, but the number of simulations required for accurate parameter approximation is inefficient for parameter space exploration. Alternatively, application of matrix methods to a truncated state space, an approach called finite state projection (FSP) [17, 18], has been applied for inference [4, 6, 13, 19] but is extremely inefficient when the state space is much larger than the number of data points. In certain cases, CMEs can be solved using generating functions that transform infinite-dimensional CMEs to complex partial differential equations (PDEs). However, this approach requires performing an inverse Fast Fourier Transform over a finite, truncated array [20]. Both FSP and generating function methods require state-space truncation, which leads to error in calculated probabilities, with a trade-off between higher accuracy and increased evaluation time with increased grid size. Further, these methods are not amenable for efficient calculation of likelihood gradients necessary for widely used machine-learning based numerical inference procedures [21, 22]. In addition, the challenge of approximation suffers from the so-called “curse of dimensionality”: the size of the state-space grows exponentially with the number of modeled species.

In the fields of machine learning and data-driven modeling, recent work has demonstrated the ability of sparse function approximations to distributions [23] and learned mappings between function spaces rather than between finite-dimensional Euclidean spaces [24] to learn generalizable solutions that govern the dynamics of complex systems. For example, [24] has introduced a neural operator formulation that models fluid dynamics in turbulent flows governed by such PDEs as the Navier-Stokes equations several orders of magnitude faster than traditional solvers.

Inspired by these approaches, we present a computationally efficient approach that uses neural networks coupled to statistical analysis of system distributions to approximate multidimensional solutions to CME models of transcriptional dynamics. In particular, we investigate a CME system involving transcriptional bursting, splicing and degradation (Figure 1a). This system of bursty transcription, although experimentally validated [25], does not yield a closed-form steady-state solution and requires costly numerical evaluation procedures [26]. We alleviate this problem by using features of the system’s univariate conditional distributions to define a statistical approximation for conditional nascent RNA distributions in terms of a mixture of negative binomial kernel functions. The weights for these kernel functions are learned by a neural network (Figure 1b). We then multiply by marginal mature RNA probability to reconstruct full joint distributions (Figure 1c). We denote this procedure kernel weight regression (KWR).

**Figure 1.**
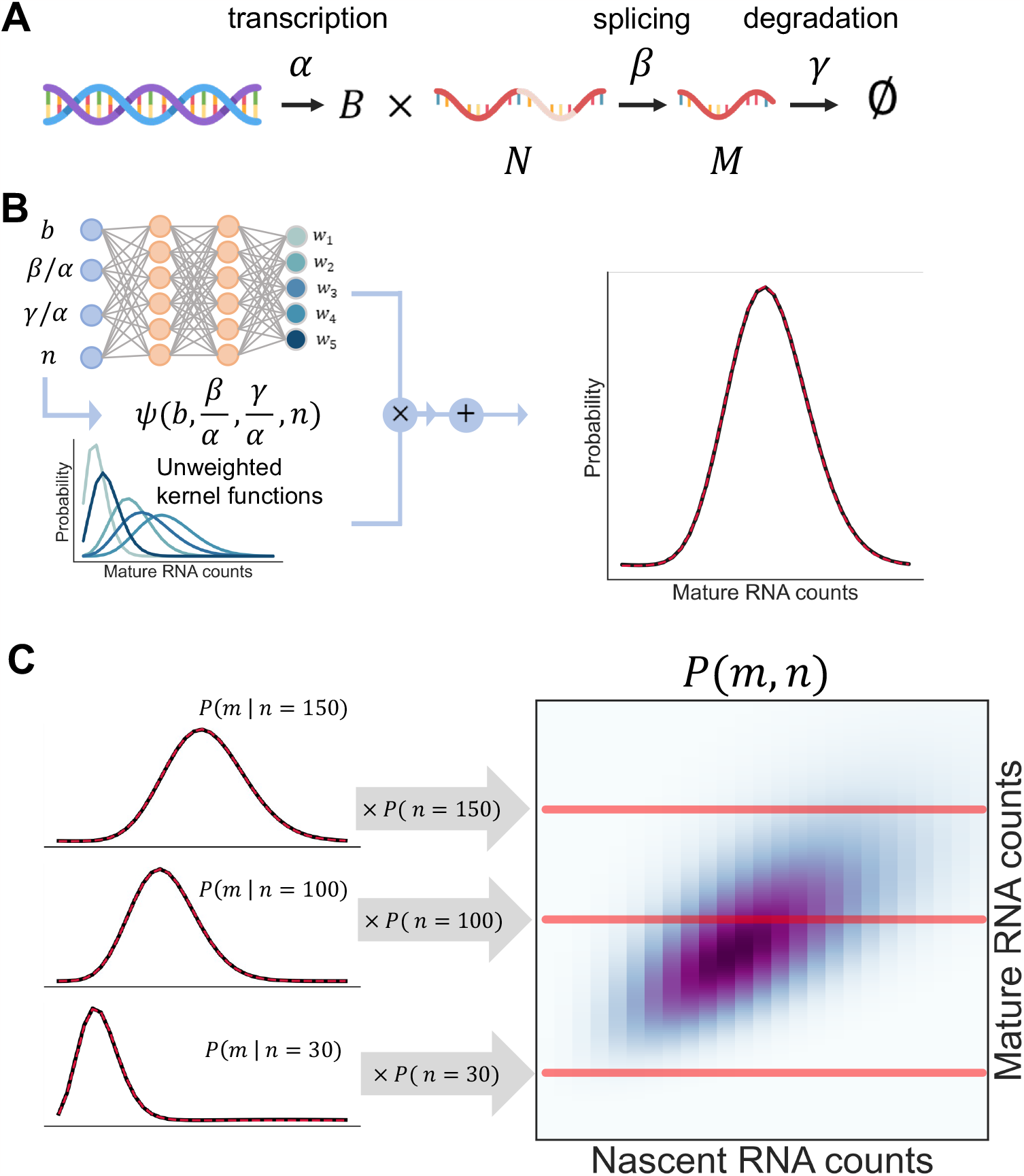
A schematic representation of the KWR approximation procedure. **A**. The biological process under consideration: bursty transcription of nascent RNA, followed by splicing to produce mature RNA and mRNA degradation. **B**. Univariate conditional distributions are approximated by summing a set of kernel functions with neural network learned weights. **C**. Joint distributions are reconstructed by multiplying conditional mature RNA probabilities by marginal nascent RNA probabilities.

To increase flexibility of the neural approximation while further decreasing evaluation time, we also implement an approximation approach by learning scaling factors for pre-calculated mean and variances of fewer negative binomial distribution kernels in addition to weights. The joint probability is again approximated by applying conditional probability. We call this approach parameter scaled kernel weight regression (psKWR). Both procedures reconstruct joint distributions by approximating conditional distributions for mature RNA molecules with neural-network scaled kernel distributions, then reconstructing full joint distributions using an analytical solution for marginal nascent RNA distribution. KWR and psKWR also provide approximations on an unbounded support and are compatible with gradient calculations required for machine-learning based likelihood optimization strategies, thereby facilitating inference of system parameters at increased resolution [27].

Our work complements other recent reports that leverage neural networks to approximate CME solutions for single-cell biophysics models. [28] use recurrent neural networks to directly learn CME solutions over a grid but cannot extrapolate beyond a predefined state space. [29] have proposed neural approximations for continuous-valued statistics of CME systems but cannot be used to perform inference. Most pertinently, [30] have independently proposed a method for approximating CME solutions with a set of negative binomial mixture functions, implemented using the Nessie framework. However, Nessie learns mixture function locations and weights, while our approach simplifies the learning process by learning weights and location scaling factors, initially placing kernel functions by exploiting the unimodality and moments of the system’s CME solution. Further, Nessie is designed to fit univariate distributions; we present a strategy for learning multivariate distributions, an essential feature for the analysis of increasingly available multimodal data.

## 2 Methods

### 2.1 Transcriptional system

We consider the system illustrated in Figure 1a, which encodes a standard model of the mRNA life-cycle [26]. Transcriptional events arrive according to a Poisson process with rate *α*, and generate *B* nascent mRNAs (*N*), where *B* is a geometric distribution with mean *b*. After an exponentially-distributed delay with rate *β*, nascent mRNAs are spliced into mature mRNAs (*M*). Finally, after an exponentially-distributed delay with rate *γ*, mature mRNAs are degraded. This system induces a stationary distribution with the probability mass function (PMF) *P* (*n, m*; *b, β, γ*), where *β* and *γ* are considered in units of *α* with no loss of generality at steady state. Although the system does not afford a closed-form solution, its joint PMF can be approximated to high precision through generating functions evaluated over a state-space grid (the larger the state-space, the greater the accuracy) [26]. Further, its nascent marginal *P* (*n*) is negative binomial [26].

### 2.2 KWR approximation overview

To approximate full joint distributions, we train a neural network to approximate the distributions of mature RNA molecules conditional on a nascent copy number *n* (Figure 1b), i.e., the univariate PMFs *P* (*m*|*n*; *b, β, γ*). We first generate rate vectors *θ*:= {*b, β, γ*} and compute high-quality generating function solutions over state-space grids to obtain joint PMFs *P* (*m, n*; *θ*), following Equation 18. These solutions are denoted QV20 (as they are evaluated on grids with limits 20 standard deviations above nascent and mature marginal means) and treated as ground truth. We then divide by nascent probabilities *P* (*n*; *θ*) to extract conditional mature probabilities *P* (*m*|*n*; *θ*).

To construct a basis for the conditional distribution at the nascent copy number *n*, we use a deterministic function *ψ* (*θ, n*) to calculate kernel means *μ*_*k*_ and standard deviations *σ*_*k*_ of negative binomial basis kernels *P*_k_, where *k* indexes the negative binomial kernels (Section S1.3). The function takes advantage of the marginal means and variances of the system, constructing an initial basis by moment-matching to an analytically tractable lognormal distribution (Section S1.3). Next, we use a neural network to map 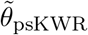, a transformation of *θ* and *n*, to a set of weights **w** and a kernel width scaling factor *h*. We train the network to minimize truncated Kullback-Leibler divergence (KL divergence, Section S1.4) between the conditional true distributions *P* (*m*|*n*; *θ*) and the conditional kernel approximations *Q*_KWR_(*m*|*n, θ*):

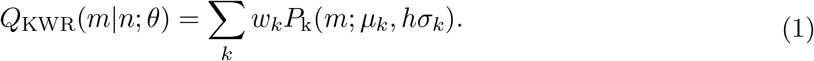

Finally, to reconstruct the joint distributions, we multiply each *Q*_KWR_(*m*|*n*; *θ*) by the analytical nascent marginal probability *P* (*n*; *θ*), as in Figure 1c:

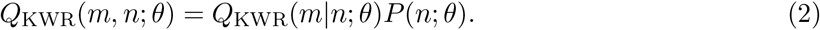

We call this method kernel weight regression (KWR), with a full description in Section S1.7. Model architecture choice is motivated in Figure S3. Figure S1 depicts final model architecture and TableS3 lists final model parameters.

### 2.3 Parameter scaled kernel weight approximation overview

To further allow for flexibility of distribution approximation, we present an alternative strategy, which scales the pre-calculated kernel means and variances. Here, we use the same function *ψ* (*θ, n*) for a given *n* to estimate kernel distribution means *μ*_*k*_ and standard deviations *σ*_*k*_. We then use a neural network to map 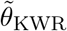, a slightly modified transformation of *θ*and *n*, to weights **w** and scaling factors **c**_*μ*_ and **c** _***σ***_ to construct the approximate conditional distribution *Q*_psKWR_(*m*|*n*; *θ*):

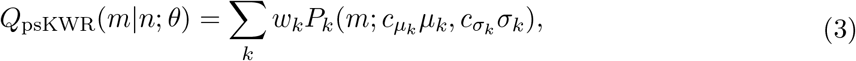

where *k* again indexes the kernel. The joint probabilities are then reconstructed by multiplying each *Q*_psKWR_(*m*|*n*; *θ*) by marginal *P* (*n*; *θ*):

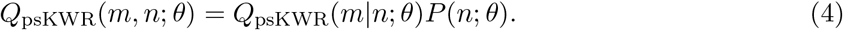

We train the network to minimize the Hellinger distance between the joint true distributions *P* (*m, n*; *θ*) and the joint approximations *Q*_*psKW R*_(*m, n*; *θ*).

We call this approach parameter scaled kernel weight regression (psKWR). It is fully described in Section S1.8, with model architecture choice motivated in Figure S4, final model architecture illustrated in Figure S2, and final model parameters listed in Table S4.

### 2.4 Data generation and generating function solutions

To generate training, validation, and testing distributions, we sampled vectors of biophysical rate vectors *θ*within the following biologically motivated bounds:

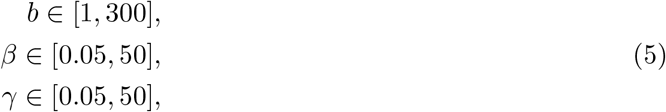

where the sampling measure was log-uniform. As described previously, *β* and *γ* are unitless quantities, scaled to the value of *α*. We generated a vector of three random numbers within the bounds. Next, we computed *μ*_*N*_ and *μ*_*M*_, the average nascent and mature copy numbers (Table S1). To ensure that training distributions matched the low-copy number regime of experimental scRNA-seq data, we restricted to vectors with *μ*_*N*_ and *μ*_*M*_ both less than 1,000. For each *θ*, we evaluated the previously derived generating function solution to the system, given in Equation 18, up to the bounds *μ*_*z*_ + 20 *σ*_*z*_ (where *z* is either nascent *n* or mature *m* value) using adaptive quadrature (implemented in the `scipy` function `integrate.quad_vec` [31]) and the inverse real fast Fourier transform (implemented in the `scipy` function `fft.irfft2` [32]). We denote this high-precision procedure by QV20 and treat it as ground truth.

We further generated lower-order generating function solutions for benchmarking. We calculated generating function solutions using adaptive quadrature on grids of 10, 4, and 1 standard deviations above nascent and mature means, or within the bounds *μ*_*z*_ + 10 *σ*_*z*_, *μ*_*z*_ + 4 *σ*_*z*_, and *μ*_*z*_ + *σ*_*z*_ in each dimension *z* ∈ {*N, M*}. We refer to these solutions as QV10, QV4, and QV1 in the text and Figure 2.

**Figure 2.**
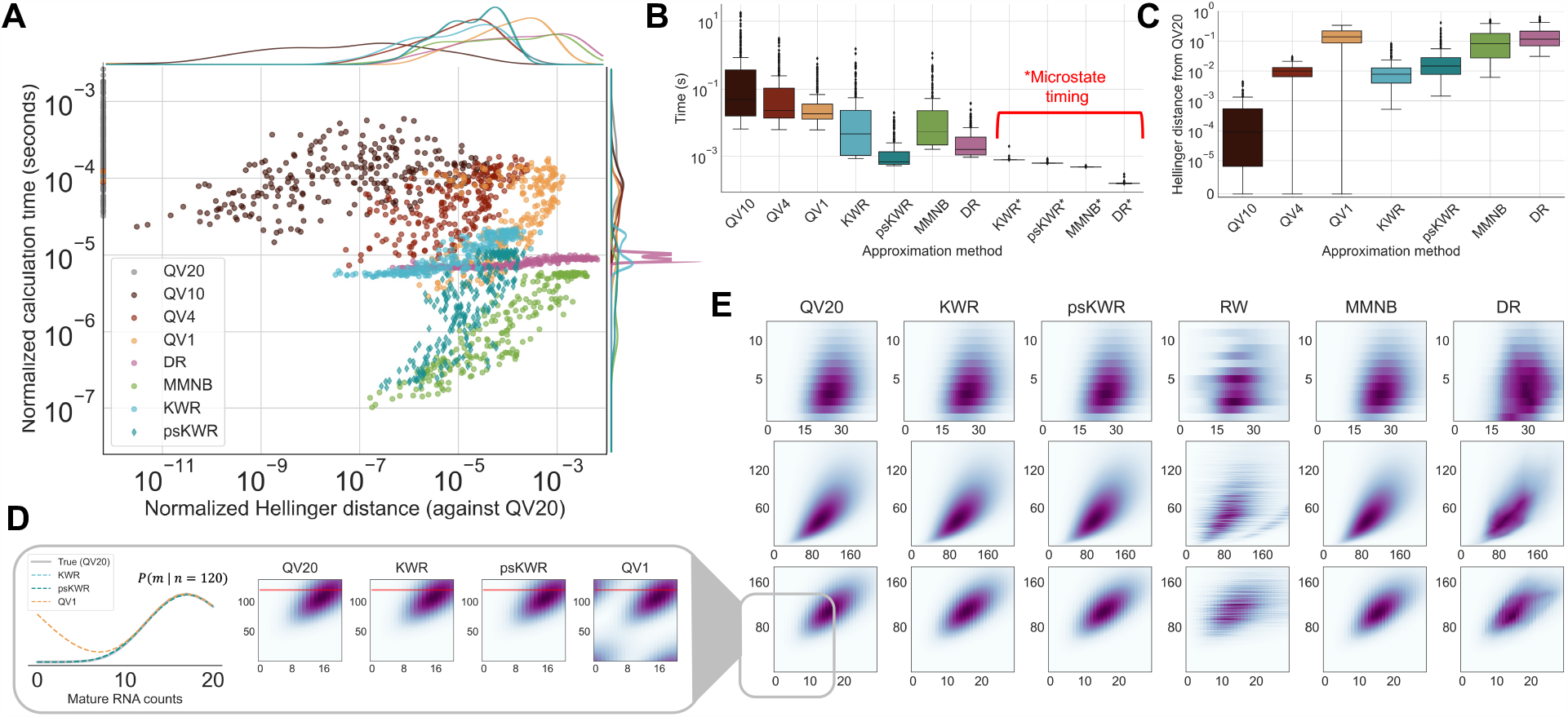
KWR and psKWR approximations accurately reconstruct joint probability distributions, with low runtime. **A**. Time to generate joint distributions over grid versus Hellinger distance, both normalized by grid size, for 256 testing rate vectors, comparing three generating function based methods (QV10, QV4, and QV1), direct regression (DR), a moment-matched negative binomial (MMNB) approximation, kernel weight regression (KWR), and parameter scaled kernel weight regression (psKWR) to ground truth (QV20). **B**. Time (unnormalized) to calculate probability for 768 testing rate vectors, where the first seven bars are grid-evaluation and an asterisk indicates method runtime for a single microstate probability. **C**. Hellinger distance (unnormalized) over grid for 768 testing rate vectors. **D**. Conditional probabilities and joint probabilities calculated using KWR, psKWR, and QV1 on a grid one standard deviation above nascent and mature means for a single testing rate vector show that significant error (as compared to QV20) is introduced on a small grid by truncation for quadrature methods but not neural approximation methods. **E**. Three example distributions approximated by KWR, psKWR, and several controls (RW, MMNB, and DR) demonstrate qualitative agreement between neural approximation methods and high-quality numerical solutions (QV20) but not all controls.

We trained KWR models on normalized conditional values and psKWR models on joint microstate probabilities. To generate normalized conditional training data for KWR, we extracted the entries corresponding to each *n* and divided them by the analytical solution for the marginal *P* (*n*; *θ*) (Equation 7, implemented in the `scipy` function `stats.nbinom.pmf` [31] with *r* = *β*^*−*1^ and *p* = (1 + *b*)^*−*1^ [33]). The resulting conditional distributions were stored along with their values of *θ* and *n*.

### 2.5 Model training and architectures

KWR models were trained to minimize the truncated KL divergence (see Section S1.4) between approximated conditional distributions *Q*_KWR_(*m*|*n*; *θ*) and normalized QV20 conditional distributions *P* (*m*|*n*; *θ*) for mature RNA transcripts.

Architectures of varying number of layers, nodes per layer, and number of kernel functions were trained and compared, with testing results for different models shown in Figure S3. The architecture chosen for final comparisons is a MLP comprising 2 hidden layers of 256 nodes the output of which is passed to one layer of 256 nodes that produces weights for 10 basis functions, and a different layer of 256 nodes that produces the scaling factor *h* (described in 2.2). This final model was trained for 35 training epochs with a learning rate of 0.001 with 1,536 training rate vectors θ (corresponding to 211,515 vectors 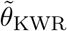 conditional vectors) and 512 validation rate vectors θ (corresponding to 72,906 vectors 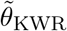 conditional vectors). All models were implemented in PyTorch with the Adam optimizer [34]. Full descriptions of training and architectures are included in Section S1.7, Figure S1, and Table S3.

psKWR models were trained to minimize the Hellinger distance between joint approximations *Q*_*psKW R*_(*m, n*; *θ*) and QV20 joint microstate probabilities *P* (*m, n*; *θ*). We again compared model architectures of different numbers of layers, nodes per layers, and basis functions to obtain a final S1.7 model (Section and Figure S3). The final model chosen for psKWR approximation is a MLP with 3 hidden layers (L1, L2, and L3) of 512 nodes per layer that produce weights **w** for five kernel functions functions. It further includes one layer of 512 nodes that takes outputs from L2 to produce scaling factors **c**_*μ*_ for kernel means described in Section layer of 512 nodes that takes outputs from L2 to produce scaling factors **c** _***σ***_ for kernel standard deviations. It was trained for 100 epochs on 1,280 rate vectors *θ*(corresponding to 211,515 conditional vectors 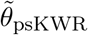)and 512 validation rate vectors *θ*(corresponding to 72,906 conditional vectors 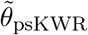). The model was implemented in PyTorch and optimized using the Adam optimizer with a learning rate of 0.001 [34]. A full S1.8 description of training is included in model parameters listed with final model architecture diagrammed in S2 and final in Table S4.

### 2.6 Inference of biophysical parameters in mouse brain cells

We implemented psKWR and KWR neural approximation strategies into an existing maximum likelihood inference framework *Monod* [35] to infer burst sizes and rates of splicing and degradation for 200 genes in Gabaergic cells in the mouse brain [36]. We obtained data from mouse sample C01 (donor ID 427378) from the Allen Institute Mouse Primary Motor Cortex (MOp) dataset [36]. Raw FASTQ files for the data were obtained and processed using `kallisto — bustools` as described in [35]. Briefly, raw reads were aligned to a reference of unspliced and spliced transcripts (generated using the flag `--lamanno)` to obtain counts for unspliced (intron containing, treated as nascent transcripts) and spliced transcripts (non-intron containing, treated as mature transcripts). Cells were filtered to remove low-quality cells (as in [35] by removing those that did not pass `kallisto — bustools` filtering and those with fewer that 10^4^ unique molecular identifier counts.

We then restricted our analysis to cells of a single cell sub-type, Gabaergic neurons, according to cell type annotations from the original study [36]. For these 1,704 subset Gabaergic cells, gene selection was performed using *Monod* to obtain genes with unspliced and spliced means (across all cells) of at least 0.01, maximum of unspliced and spliced counts (across all cells) of at least 3, and maximum spliced counts (across all cells) no greater than 400. We then selected the top 200 of remaining genes by expression (as in [35]).

For all 200 genes, inference was performed to obtain burst size *b*, relative splicing rate *β/α*, and relative degradation rate *γ/α*, according to the bursty model of transcription (Section 2.1), by maximizing data likelihood under the model’s steady-state distribution. For all genes, three different methods of evaluating data likelihoods were used: 1) the default generating function method with adaptive quadrature (QV, Section S1.2, 2) KWR approximation, and 3) psKWR approximation. Inference bounds of [*−*1.0, 4.2], [*−*1.8, 2.5],[*−*1.8, 3.5] were set for log_10_*b*, log_10_*β*, and log_10_*γ*, respectively. Inference was performed with a maximum of 15 iterations, a single search (no restarts), and parameters initiated at their method of moments estimates.

## 3 Results and discussion

To benchmark the algorithm’s performance, we investigated the accuracy and runtime for PMF grid evaluation for a set of testing rate vectors (Figure 2a). To quantify accuracy, we calculated the Hellinger distance, which measures the discrepancy between probability distributions *P* and *Q* and takes the form 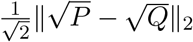, using the high-quality QV20 solutions as ground truth.

### 3.1 Comparison to generating function grid evaluations

We first compared KWR to three lower-order generating function-based solutions, denoted QV1, QV4, and QV10, which evaluate the numerical steady-state solution (Section S1.6) to varying accuracies based on grid size. We found that the KWR approximation achieves accuracy comparable to lower-order generating function solutions while reducing evaluation time (Figure 2a). For example, the average Hellinger distances for testing examples (normalized by grid size) were 3.085 × 10^*−*5^ and 3.370 × 10^*−*6^ for QV4 and KWR, respectively, but KWR approximation was more than 6 times faster (even over a grid) with an average time of 9.6307 × 10^*−*6^ seconds for QV4 evaluation and 1.624 × 10^*−*6^ seconds for KWR approximation (Figure 2**A**). KWR was both faster and almost an order of magnitude more accurate than generating function solution QV1, which had the larger average evaluation time (normalized by grid size) of 4.978 × 10^*−*5^ seconds and the greater Hellinger distance of 2.637 × 10^*−*4^ (both normalized by grid size, Figure 2**A**). While psKWR was slightly less accurate than KWR, its accuracy was still comparable to the generating function solutions ofQV4 and more accurate than QV1 with a 3.976 × 10^*−*5^ average Hellinger distance. psKWR even further reduced evaluation time with an average normalized grid evaluation runtime of 4.414 × 10^*−*6^ seconds, two orders of magnitudes faster than QV1 (Figure 2**A**). This benchmark used 256 testing examples, with Hellinger distances and runtimes normalized by the grid size.

#### Comparison to statistical controls and baseline models

As a control to assess how learning kernel function weights contributes to reconstruction accuracy, we investigated approximations obtained using random weights (RW) **w** in Equation 1. RW employs the same function *ψ*_*KWR*_(*θ, n*) to generate kernel means and variances as KWR, and identically scales variances by learned factor *h*, but weights kernels with random weights that sum to unity. As shown quantitatively in Figure 2**B**, RW does not accurately reconstruct joint probabilities, motivating the use of learned weights.

To explore how much accuracy improves when using a neural network to scale means and variances rather than using mere statistical approximations, we also benchmarked against a non-neural, statistical approximation (moment-matched negative binomial or MMNB). The MMNB approach (Section S3) generates statistically plausible means and variances for mature RNA conditional distributions (using the same function *ψ*_*KWR*_(*θ, n*)). These means and variances parameterize a single negative binomial distribution, which are then multiplied by the analytical nascent marginal probability to produce join probability approximations. MMNB underperforms KWR and psKWR in Hellinger distance accuracy over a grid, evidence that the neural procedure is necessary to achieve highly accurate approximations.

To assess the utility of kernel functions, we also compared KWR and psKWR to a method of direct approximation that uses a fully connected multilayer perceptron to map from an input *{θ, n, m}* to a single-valued joint probability *P* (*m, n*; *θ*). The network was trained to reduce the mean squared error between numerically calculated and predicted joint probabilities. We refer to this method as direct regression (DR), described in Section S2 with testing results in Figure S5 and network parameters in Table S5. DR was considerably less accurate than the KWR and psKWR approximations (Figure 2**A,B**). This highlights that basis function learning methods like KWR and psKWR, which output parametric forms rather than single probabilities, are desirable and efficient for this application.

#### Evaluation time for microstate probabilities

In the context of statistical inference from data, optimizing likelihoods involves the evaluation of a single microstate *P* (*n, m*; *θ*). Here, KWR and psKWR outperform generating function-based methods by design by bypassing grid evaluation. Generating function evaluation over grids can take tens of seconds (QV10, Figure 2**B**), extremely prohibitive for inference processes in high-throughput genomics in which likelihoods must be iteratively calculated for thousands of genes. Evaluating single microstate probabilities using the neural approximations, by contrast, took 4.205 × 10^*−*2^ and 3.284 × 10^*−*3^ seconds on average, decreasing evaluation time by three and four orders of magnitude (Figure 2**B**) with similar accuracies (Figure 2**A**,**C**). While DR and MMNB slightly outperform KWR on microstate timing (Figure 2**B**), their accuracy is substantially lower (Figure 2**A**,**C**). Further, psKWR has evaluation time comparable to DR and MMNB and is more accurate (Figure 2**A**,**B**,**C**).

#### Joint distribution reconstruction

Some examples of probability distributions obtained by the approximation procedures are shown in Figure 2**D**, where we again treat QV20 as ground truth. Both KWR and psKWR approximations are qualitatively indistinguishable from ground truth, suggesting that they are appropriate for moderate-precision inference. While MMNB results look qualitatively similar, Hellinger distances from ground truth are substantially greater (Figure 2**A**,**C**), suggesting that while it is appropriate as a coarse approximation, it may not be not sufficient for precise inference. The random weight control recapitulates the general shape of the distribution, but introduces considerable error into microstate probabilities, confirming that the learned weights are essential for an accurate approximation (Figure 2**D**, fourth column). Finally, the DR results noticeably distort the distribution shapes, further evidencing the utility of the kernel function approach (Figure 2**D**, sixth column).

#### KWR and psKWR use in biophysical parameter inference

KWR and psKWR can be used in methods in an existing maximum likelihood framework [35] to infer biophysical parameters from experimental data. We used both neural approximation strategies and the default generating function solver to infer burst sizes (denoted QV in Figure 3), relative splicing rates, and relative degradation rates for 200 highly expressed genes in Gabaergic cells from the mouse primary motor cortex (see Section 2.6) [36].

**Figure 3.**
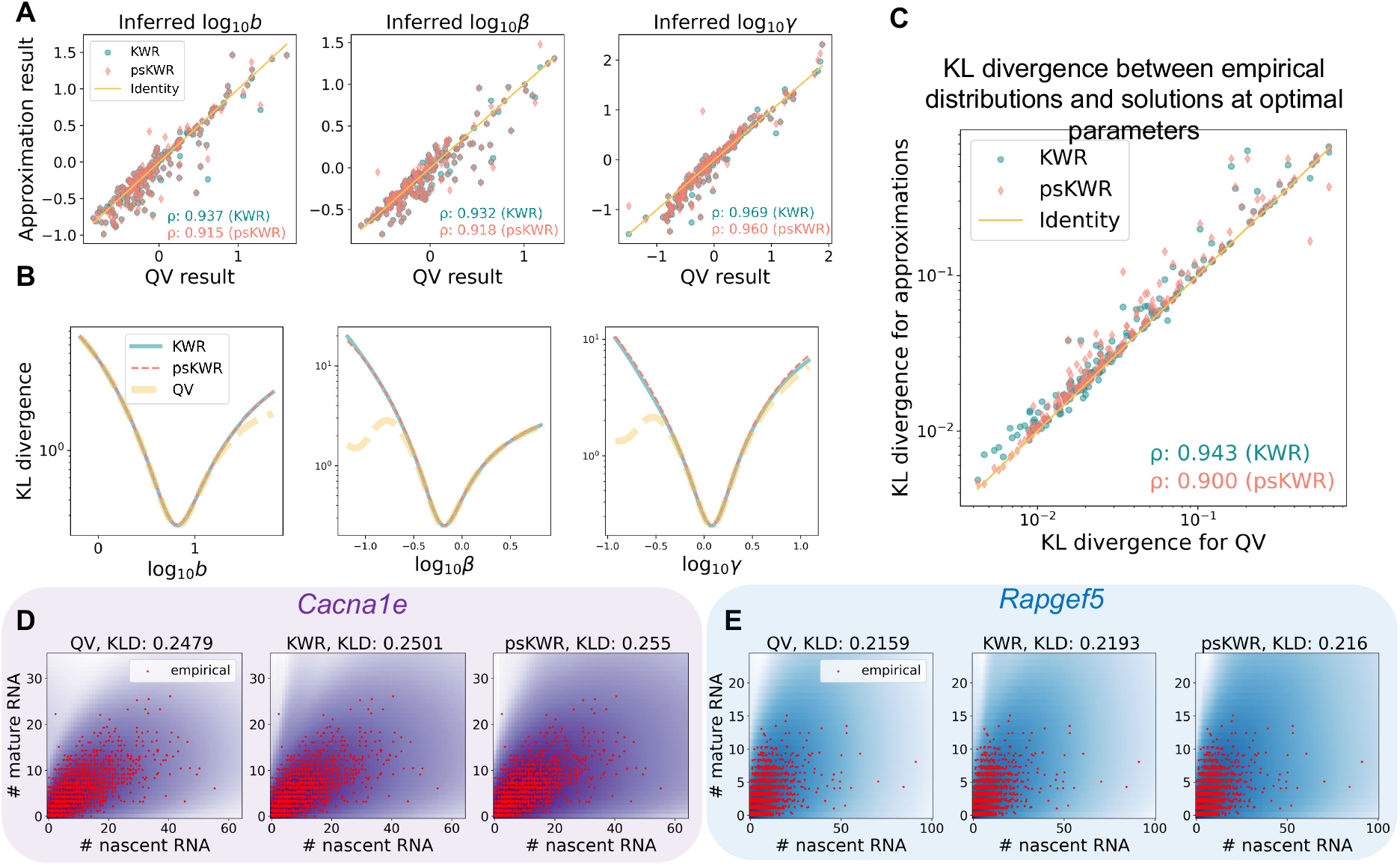
Use of KWR and psKWR in inference frameworks results in likelihoods and parameters consistent with those obtained using standard numerical solutions, with better behaved likelihood landscapes. **A**. Parameters for the bursty model of transcription (burst size *b*, splicing rate *β*, and degradation rate *γ*, with splicing and degradation rate considered in units of transcription rate *α*) highly correlate with results using adaptive quadrature. **B**. For the gene *Cacna1e*, Kullback-Leibler divergence between data and biophysical model likelihoods calculated at two of the optimal parameters with the indicated third varied. Likelihood landscapes for biophysical parameters evaluated using KWR and psKWR show one clear local and global minimum, while truncation necessary for numerical solution QV introduces off-optimum decreases in KL divergence. **C**. The log_10_ of lowest KL divergence obtained between data distributions and neural approximation strategies and data distributions and QV solutions at their respective optimal parameters highly correlate. **D**. Joint count distributions for *Cacna1e* and **E**. *Rapgef5*, both empirical (red scatter points) and evaluated using QV, KWR, and psKWR (colored density) at the optimal parameters they respectively produced.

For the 200 fit genes, we found that inference results using KWR and psKWR approaches for data likelihood evaluation highly correlate with results using the standard generating function approach (Figure 3**A**), with better defined likelihood landscapes (Figure 3**B**). Pearson correlations of greater than 0.9 were obtained for log_10_ of inferred burst sizes, relative splicing rates, and relative degradation rates for KWR against QV and psKWR against QV (Figure 3**A**). KL divergence between empirical and model distributions at optimal inferred parameters also agree between QV and both neural approximation strategies (Figure 3**C**). In addition, distributions produced by all methods capture empirical RNA distributions, with two example genes shown in Figure 3**D** and **E**. These results suggest that KWR and psKWR are reliable alternatives to numerical solutions for parameter inference.

## Conclusion

After training on a relatively small set of example rate vectors, the KWR and psKWR neural approximation approaches generalize to produce highly accurate and unbounded solutions to a mechanistic model of the lifecycle of RNA in single cells (Figure 2). They are as accurate as lower-order generating function methods, but bypass quadrature and Fourier inversion, achieving orders of magnitude faster runtimes. Thus, these approaches can greatly reduce the amount of time necessary to explore steady-state distributions across a wide range of transcriptional parameters for inference, parameter sensitivity investigations, and experimental design [37]. In particular, we have recently employed KWR application in a popular variational autoencoder framework to infer biophysical parameters for thousands of cells and genes, made possible because KWR, unlike generating function solutions, is compatible with gradient descent algorithms 21,27.

Recent advances in simultaneous quantification [38] and modeling [39] of mRNA and proteins present natural further applications as trivariate unspliced mRNA – spliced mRNA – protein data are available, but the corresponding stochastic system does not afford an analytical solution. Single cell measurements of accessible regions of chromatin, obtained using ATAC-seq [40], may also be modeled using CMEs to produce multivariate steady-state distributions, an example of yet another system for which our neural approximation strategies can facilitate mechanistic analysis. As single-cell assays with multiple molecular species become increasingly more common, so increases the need for computationally efficient methods for modeling and inference that go beyond statistical summaries to yield mechanistic insight into system dynamics.

## Supporting information

spectral_neural_supplement

## 4 Data and code availability

All training and validation datasets, trained models, and scripts to generate manuscript figures are available at https://github.com/pachterlab/GCCP_2022.

## 5 Author contributions

G.G. conceptualized the neural approximation strategies and was involved in implementation, statistical analyses, and basis function design. M.C. implemented and adapted the strategies, trained models, and performed analyses. T.C. and L.P. provided feedback, design suggestions, and trouble-shooting advice. All authors wrote the article.

## 7 Acknowledgments

We thank Dr. Yisong Yue, Yongin Choi, Dr. John J. Vastola, and Dr. Zachary Fox for valuable discussions. G.G. and L.P. were partially funded by NIH U19MH114830. M.C. is funded by the National Science Foundation Graduate Research Fellowship Program and Amazon AI4Science Fellowship. Figures 1, 2, and S3 use color palettes from MetBrewer, developed by Blake R. Mills and available at https://github.com/BlakeRMills/MetBrewer.

