## Supplementary material for "Spectral neural approximations for models of transcriptional dynamics": spectral_neural_supplement

### Spectral neural approximations for models of transcriptional dynamics
### Supplementary Material

Gennady Gorin[1,*], Maria Carilli[1,2,*], Tara Chari[2], and Lior Pachter[2,3,**]

[1]Division of Chemistry and Chemical Engineering, California Institute of Technology, Pasadena, CA, 91125
[2]Division of Biology and Biological Engineering, California Institute of Technology, Pasadena, CA, 91125
[3]Department of Computing and Mathematical Sciences, California Institute of Technology, Pasadena, CA, 91125
[*]These authors contributed equally to this work.

November 30, 2023

### S1 Supplementary Methods

#### S1.1 Statistical preliminaries

The Poisson distribution over the non-negative integers $x \in \mathbb{N}_0$ is parameterized by its mean $\lambda$:

$$P_{Poiss}(x; \lambda) = \frac{\lambda^x e^{-\lambda}}{x!}. \tag{6}$$

The negative binomial (NB) distribution over $x \in \mathbb{N}_0$ can be parameterized by the shape $r$ and the success probability $p$:

$$P_{NB,rp}(x; r, p) = \frac{\Gamma(r + x)}{x!\Gamma(r)}(1 - p)^x p^r. \tag{7}$$

This is the probability mass function (PMF) in its combinatorial form, which follows the convention used in MATLAB, `numpy`, and `scipy`.

Alternatively, the NB distribution can be parametrized by the shape $r$ and the mean $\mu$:

$$P_{NB,r\mu}(x; r, \mu) = \frac{\Gamma(r + x)}{x!\Gamma(r)} \left(\frac{r}{r + \mu}\right)^r \left(\frac{\mu}{r + \mu}\right)^x. \tag{8}$$

The shape $r$ is specified given a mean and standard deviation (as long as $\sigma^2 > \mu$):

$$r(\mu, \sigma) = \frac{\mu^2}{\sigma^2 - \mu}. \tag{9}$$

We use the resultant mean-variance $(\mu, \sigma^2)$ parametrization of the NB distribution:

$$P_{NB}(x; \mu, \sigma) = P_{NB,r\mu}(r(\mu, \sigma), \mu). \tag{10}$$

The lognormal distribution over $x \in \mathbb{R}_+$ can be parametrized by the mean and standard deviation of the underlying exponentiated normal distribution (from here, referred to as the log-mean $\mu$ and the log-standard deviation $\sigma$):

$$f(x; \mu_l, \sigma_l) = \frac{1}{x\sigma_l\sqrt{2\pi}} \exp\left(-\frac{(\ln x - \mu_l)^2}{2\sigma_l^2}\right). \tag{11}$$

The parameters can be set according to the distribution's desired mean and variance:

$$\begin{aligned} \mu_l &= \ln \frac{\mu^2}{\sqrt{\sigma^2 + \mu^2}}, \\ \sigma_l &= \left[\ln \frac{\sigma^2 + \mu^2}{\mu^2}\right]^{1/2}. \end{aligned} \tag{12}$$

The quantiles of the lognormal distribution have the following form:

$$q(x; \mu_l, \sigma_l) = \exp\left(\mu_l + \sigma_l \Phi^{-1}(x)\right), \tag{13}$$

where $\Phi$ is the cumulative distribution function of the standard normal distribution and $\Phi^{-1}$ is its quantile function.

The bivariate lognormal distribution over $x, y \in \mathbb{R}_+$ can be similarly parametrized, with an additional parameter that governs the correlation of the underlying bivariate normal distribution (from here, referred to as the log-correlation $\rho_l$):

$$
\begin{aligned}
f(x, y; \mu_{xl}, \mu_{yl}, \sigma_{1l}, \sigma_{yl}, \rho_l) &= \frac{1}{xy \times 2\pi\sigma_{1l}\sigma_{yl}\sqrt{1-\rho_l^2}} \exp\left(-\frac{A^2 + B^2 - 2\rho_l AB}{2(1-\rho_l^2)}\right), \\
A &= \frac{\ln x - \mu_{1l}}{\sigma_{1l}}, \\
B &= \frac{\ln y - \mu_{yl}}{\sigma_{yl}},
\end{aligned}
\tag{14}
$$

where $\mu_{xl}$ and $\mu_{yl}$ are the marginal log-means, $\sigma_{xl}$ and $\sigma_{yl}$ are the marginal log-standard deviations [41]. The log-correlation $\rho_l$ can be set according to the distribution's desired marginal moments and correlation $\rho$:

$$
\rho_l = \frac{1}{\sigma_{xl}\sigma_{yl}} \ln\left(\rho\sigma_1\sigma_2 e^{\mu_{xl}+\mu_{yl}+(\sigma_{xl}+\sigma_{yl})/2} + 1\right). \tag{15}
$$

Finally, the conditional distribution of $y$ given $x$ is lognormal, with the following log-mean and log-standard deviation:

$$
\begin{aligned}
\mu_{yl,x} &= \mu_{yl} + \rho_l \frac{\sigma_{yl}}{\sigma_{xl}}(\ln x - \mu_{yl}), \\
\sigma_{yl,x} &= \sigma_{yl}\sqrt{1-\rho_l^2}.
\end{aligned}
\tag{16}
$$

#### S1.2 System definition

We consider the following system:

$$
\varnothing \xrightarrow{\alpha} B \times \mathcal{N} \xrightarrow{\beta} \mathcal{M} \xrightarrow{\gamma} \varnothing, \tag{17}
$$

where $\mathcal{N}$ denotes a nascent mRNA transcript and $\mathcal{M}$ denotes a mature mRNA transcript (Figure 1**A**). The system's rates and burst size are defined in Table S1. We set the burst frequency $\alpha$ to unity and consider all other rates in units of $\alpha$ with no loss of generality at equilibrium.

The joint generating function solution was derived in [26]:

$$
\begin{aligned}
G(x_N, x_M) &= \sum_{x_N, x_M = 0}^{\infty} x_N^n x_M^m P(n, m) \\
u_z &= x_z - 1 \\
U(u_N, u_M, s) &= u_M \frac{\beta}{\beta - \gamma} e^{-\gamma s} + \left(u_N - u_M \frac{\beta}{\beta-\gamma}\right) e^{-\beta s} \\
\ln G := \phi(u_N, u_M) &= \int_0^\infty \left[\frac{1}{1 - bU(u_N, u_M, s)} - 1\right] ds.
\end{aligned}
\tag{18}
$$

| Symbol | Definition |
|---|---|
| $N$ | Nascent mRNA species index |
| $M$ | Mature mRNA species index |
| $z$ | Generic species subscript ($N$ or $M$) |
| $n$ | Nascent mRNA copy number |
| $m$ | Mature mRNA copy number |
| $P(n,m)$ | True bivariate PMF of nascent and mature RNA counts |
| $B$ | Geometric burst size distribution defined on $\mathbb{N}_0$ |
| $b$ | Mean burst size |
| $\beta$ | Splicing rate, normalized to burst frequency |
| $\gamma$ | Degradation rate, normalized to burst frequency |
| $\mu_N = b/\beta$ | Mean of nascent count distribution |
| $\mu_M = b/\gamma$ | Mean of mature count distribution |
| $\mu_z$ | Mean of generic species $z$ |
| $\sigma_N^2 = \mu_N(1+b)$ | Variance of nascent count distribution |
| $\sigma_M^2 = \mu_M\left(1+\frac{b\beta}{\beta+\gamma}\right)$ | Variance of mature count distribution |
| $\sigma_z^2$ | Variance of generic species $z$ |
| $\mathrm{Cov}(N,M) = \frac{b^2}{\beta+\gamma}$ | Covariance of nascent and mature RNA counts |
| $\rho = \frac{\mathrm{Cov}(N,M)}{\sigma_N\sigma_M}$ | Pearson correlation of nascent and mature RNA counts |

Table S1: Statistical and biological quantities of interest.

Here $x_N$ and $x_M$ are complex generating function arguments. The nascent marginal of this distribution is given by a negative binomial law [26,33]:

$$P(n) = P_{NB,r\mu}\left(\frac{k}{\beta}, \frac{kb}{\beta}\right). \tag{19}$$

Salient lower moments can be computed (Table S1), and the distribution is known to be unimodal [42]. However, the microstate, conditional, and mature marginal probabilities do not have closed-form solutions.

The distribution can be evaluated on a grid of size $\mathfrak{s}_N \times \mathfrak{s}_M$, or the direct product of species-specific microstates $[0,1,...,\mathfrak{s}_N-1] \times [0,1,...,\mathfrak{s}_M-1]$, by evaluating $G$ on the complex bivariate unit sphere and performing an inverse fast Fourier transform (IFFT) [20,26]. As the IFFT sums to unity, the selected state space sizes $\{\mathfrak{s}_N, \mathfrak{s}_M\}$ must be sufficiently large to limit truncation error. Therefore, it is typically set according to standard concentration inequalities: for example, Chebyshev's inequality guarantees that the amount of probability mass at $x \geq \mu + 4\sigma$ will be at most 1/16.

#### S1.3 Overview of approximation procedures

In the most general formulation, a CME model of biology converges to a stationary distribution $P(z;\theta)$ over $z \in \mathbb{N}_0$, which can be computed at considerable computational cost through generating function inversion. This distribution may be marginal or conditional. In certain cases, $P$ affords a real Poisson representation [43]:

$$P(z;\theta) = \int_0^\infty P_{\text{Poiss}}(z;\lambda)dF_\lambda, \tag{20}$$

where $F$ is a mixing cumulative distribution function (CDF), or stationary solution to an underlying continuous-valued Fokker-Planck equation. If such a representation exists, we can always write down a finite approximation over $K$ Poisson kernels:

$$Q(z;\theta) = \sum_{k=1}^{K} w_k P_{\text{Poiss}}(z;\lambda_k), \tag{21}$$

where $w_k$ are weights on a $K$-dimensional simplex. Formally, this approximation substitutes the continuous CDF $F$ with a piecewise constant CDF $F^*$:

$$F^*(\lambda) = \sum_{k=1}^{K} w_k \mathbb{I}_{\lambda \leq \lambda_k}, \tag{22}$$

where $\mathbb{I}$ is the indicator function. In principle, every $P$ can be analyzed to produce an optimal set of weights $w_k$ and mixture components $\lambda_k$, i.e., ones which minimize a measure of statistical divergence between $P$ and $Q$. Further, as $K \to \infty$, $Q \to P$. However, the problem of actually obtaining the optimal $w_k$ and $\lambda_k$ may be challenging. More problematically, convergence in the number of kernels in $K$ is typically slow.

Our approach is to first accelerate convergence in $K$. To accomplish this, we use negative binomial kernels, which are continuous Poisson mixtures with a gamma mixing density:

$$\begin{aligned} P_{\text{ker}}(z;\mu,\sigma) &= P_{\text{NB}}(z;\mu_k,\sigma_k) \text{ if } (\sigma_k)^2 > \mu_k; \\ &= P_{\text{Poiss}}(z;\mu_k) \text{ otherwise,} \end{aligned} \tag{23}$$

where $\mu_k$ and $\sigma_k$ are the mean and standard deviation of the negative binomial kernel. Therefore, the approximation is a mixture of $K$ kernels:

$$Q(z;\theta) = \sum_{k=1}^{K} w_k P_{\text{ker}}(m;\mu_k,\sigma_k). \tag{24}$$

To implement this approximation, we need to find values of $w_k$, $\mu_k$, and $\sigma_k$. In principle, this can be accomplished by training a neural network to predict these quantities based on $\theta$; this is the approach taken in the Nessie framework [30]. However, we find that $\mu_k$ and $\sigma_k$ can be assigned based on the shape of the approximated distribution.

We seek to judiciously place the kernel locations $\mu_k$. These locations should be adaptive with respect to rate vector $\theta$, and prioritize high-probability mass regions of the distribution $P$. We assume that $P(z;\theta)$ resembles a lognormal distribution, which is typical of biomolecules [44] and consistent with the knowledge that $P$ is defined on the positive numbers, unimodal, and skewed to the right. If we have such an approximating lognormal distribution with parameters $\mu_l$ and $\sigma_l$, we can place $\mu_k$ at the quantiles of this distribution, given by Equation 13. We found that quantiles placed at the Chebyshev nodes (rescaled to $[0,1]$) gave the best performance:

$$\begin{aligned} x_k &= \frac{1}{2}\left[\cos\left(\frac{2k-1}{2K}\pi\right)+1\right], \\ \mu_k &= q(x_k;\mu_l,\sigma_l). \end{aligned} \tag{25}$$

Next, we seek to specify the kernel standard deviations $\sigma_k$. Knowing that $Q$ should be unimodal, we implement an adaptive procedure:

$$\begin{aligned} \sigma_k &:= \mu_{k+1} - \mu_k, \\ \sigma_k^* &\propto \sigma_k, \end{aligned} \tag{26}$$

i.e., the width of each kernel is controlled by the spacing between neighboring kernels. This approach prevents the individual kernels from becoming too sharply peaked. We define the constant of proportionality as $h$, such that $\sigma_k^* = h \times (\mu_{k+1} - \mu_k) := h\sigma_k$, where $\sigma_k$ is the unscaled kernel width. We set the final parameter $\sigma_k$ to $\sqrt{\mu_k}$, making the $K$th kernel Poisson.

Given this framework, we can split the problem of evaluating $Q$ into two parts. The analytical part uses the lognormal approximation to obtain the kernel locations and unscaled widths:

$$\psi : \theta, n \mapsto \begin{bmatrix} (\mu_1, \sigma_1) \\ ... \\ (\mu_k, \sigma_k) \end{bmatrix}. \tag{27}$$

As a shorthand, we use the following component-specific notation:

$$\psi_k : \theta, n \mapsto (\mu_k, \sigma_k), \tag{28}$$

with the caveat that this shorthand corresponds to extracting entries of the $K$-dimensional array in Equation 27, as $\sigma_k$ are computed in a coupled fashion.

The second part of the problem involves evaluating $\mathbf{w}$ and $h$. Conceptually, these are implemented using a neural network $\mathcal{F}$:

$$\mathcal{F} : \tilde{\theta}_{\text{KWR}} \mapsto w_1, ..., w_K, h = \mathbf{w}, h, \tag{29}$$

where $\tilde{\theta}_{\text{KWR}}$ is an transformation of $\theta$ and n.

This procedure generates a prediction $Q(z;\theta)$ for a microstate $m$ under $\theta$. By minimizing a measure of statistical distance between $P$ and $Q$, we can train the neural network $\mathcal{F}$ to output optimal $\mathbf{w}$ and $h$.

To increase flexibility and representative power of the weighted kernel functions, we present an alternative approach to approximating $P$. We note that means $\mu_k$ and standard deviations $\sigma_k$ produced by $\psi$ can be scaled by learned factors:

$$Q_{\text{psKWR}}(z;\theta) = \sum_{k=1}^{K} w_k P_{\text{ker}}(m; c_{\mu_k}\mu_k, c_{\sigma_k}\sigma_k). \tag{30}$$

This can be accomplished by training a different neural network to produce weights and scaling factors for means and standard deviations:

$$\mathcal{F}_{\text{psKWR}} : \tilde{\theta}_{\text{psKWR}} \mapsto w_1, ..., w_K, c_{\mu_1}, ..., c_{\mu_K}, c_{\sigma_1}, ..., c_{\sigma_K} = \mathbf{w}, \mathbf{c}_{\mu_k}, \mathbf{c}_{\sigma_k}, \tag{31}$$

. By minimizing a measure of statistical distance between $P$ and $Q_{\text{psKWR}}$, we can train the neural network $\mathcal{F}_{\text{psKWR}}$ to output optimal $\mathbf{w}$, $\mathbf{c}_\mu$, and $\mathbf{c}_\sigma$.

#### S1.4 Measures of statistical distance

To train $\mathcal{F}$, we use a variant of the Kullback-Leibler divergence (KL divergence) between $P$ and $Q$ as our loss function. Given a target distribution $P$ and an approximate distribution $Q$, the KL divergence is defined as follows:

$$D(P \parallel Q) = -\sum_{z=0}^{\infty} P(z) \ln \frac{Q(z)}{P(z)}. \tag{32}$$

Although this loss function is standard [30], it poses problems in the current context. Although the KL divergence is guaranteed to be positive, its truncated version (with the sum taken up to a finite limit $\mathfrak{s}$) is not. To construct a well-defined measure of error, we use the distributions' truncated analogues:

$$P'(z) := \frac{P(z)}{\sum_{z=0}^{\mathfrak{s}} P(z)}. \tag{33}$$

Therefore, the loss function takes the following form:

$$L = D(P' \parallel Q') = -\sum_{z=0}^{\mathfrak{s}} P'(z) \ln \frac{Q'(z)}{P'(z)}. \tag{34}$$

We use this version whenever we refer to the optimization of KL divergence. It is conceivable that the truncation procedure in Equation 33 can introduce error. Therefore, to benchmark the performance of the algorithm, we use an orthogonal metric, the Hellinger distance. This metric is given in Equation 35; since squares are strictly positive, the truncated version can be safely treated as a lower bound on the Hellinger distance over the entire state space.

To train $\mathcal{F}_{\text{psKWR}}$ and to quantify precision between approximated distributions for testing parameters and QV20 solutions, we calculate the Hellinger distance [30], a metric of discrepancy between probability distributions:

$$H(P, Q; \theta) = \frac{1}{\sqrt{2}} \sqrt{\sum_{n=0}^{\mathfrak{s}_N} \sum_{m=0}^{\mathfrak{s}_M} \left( \sqrt{P(n, m; \theta)} - \sqrt{Q(n, m; \theta)} \right)^2}, \tag{35}$$

where $\mathfrak{s}_N$ and $\mathfrak{s}_M$ are the state space bounds for each $\theta$.

#### S1.5 Approximating joint distributions

To approximate the joint distribution of nascent and mature RNA, we use nascent marginals and approximate mature conditionals using the approximation procedure described in Section S1.3 based on $P(n, m) = P(n)P(m|n)$. The nascent marginal $P(n)$ is known (Equation 19). To obtain the joint distribution, we need to solve the univariate problem of approximating $P(m|n)$ by a judiciously chosen $\psi$ and neural network input $\tilde{\theta}$.

To define $\psi$, we first construct a univariate lognormal distribution that resembles the true conditional distribution $P(m|n; \theta)$. As the conditional moments of $P(m, n; \theta)$ are not known in closed form, we first approximate the joint distribution $P(m, n; \theta)$ using the bivariate lognormal distribution described in Equation 15, matched to $\mu_N$, $\sigma_N^2$, $\mu_M$, $\sigma_M^2$, and $\rho$ of $P(m, n; \theta)$ (Table S1). To do so, we apply Equation 12 for $N$ and $M$ to obtain the marginal parameters, then Equation 15 to obtain the correlation. Finally, we evaluate Equation 16 at $x = n + 1$ to obtain the parameters $\mu_l$ and $\sigma_l$ of a coarse lognormal approximation of conditional distributions $P(m|n; \theta)$. The kernel locations and unscaled widths follow from Equations 25 and 26.

Next, for the kernel weight regression (KWR) approach, we define conditional examples $\tilde{\theta}_{\text{KWR}}$, transformations of $\theta, n$, to use as inputs to the network $\mathcal{F}$. We found that $\tilde{\theta}_{\text{KWR}} = \{\log_{10} b, \log_{10} \beta, \log_{10} \gamma, \mu_l, \sigma_l, \mathfrak{s}_M, n\}$, where $\mathfrak{s}_M = \mu_M + 4\sigma_M$ is the state space bound for mature counts, reduced training time as conditional means and variances were pre-calculated. Each rate vector $\theta = b, \beta, \gamma$ corresponds to $\mathfrak{s}_N$ conditional examples $\tilde{\theta}_{KWR}$. With the kernels and the network inputs specified, we iterate over pre-computed distributions $P(m|n; \theta)$ and train the network by minimizing the loss in Equation 34.

For the parameter scaled kernel weight regression (psKWR) approach, we define slightly modified conditional examples $\tilde{\theta}_{\text{psKWR}}$, transformations of $\theta, n$, to input to network $\mathcal{F}_{\text{psKWR}}$. $\tilde{\theta}_{\text{psKWR}} = \{\log_{10} b, \log_{10} \beta, \log_{10} \gamma, n, \mu_1, ..., \mu_K, \sigma_1, ..., \sigma_K\}$, where $\mathfrak{s}_M = \mu_M + 4\sigma_M$, where $\mu_k$ and $\sigma_k$ are the means and standard deviations produced by function $\psi$ as described in Section S1.3. Again, ach rate vector $\theta = b, \beta, \gamma$ corresponds to $\mathfrak{s}_N$ conditional examples $\tilde{\theta}_{psKWR}$. We then train the network by minimizing Hellinger distance between approximated and stored highly accurate numerical distributions.

#### S1.6 Data generation

To generate training and validation distributions, we sampled vectors of biophysical rate vectors $\theta$ within the following biologically motivated bounds:

$$\begin{aligned} b &\in [1, 300], \\ \beta &\in [0.05, 50], \\ \gamma &\in [0.05, 50], \end{aligned} \tag{36}$$

where the sampling measure was log-uniform. As described previously, $\beta$ and $\gamma$ are in units of $\alpha$. We generated a vector of three random numbers within the bounds. Next, we computed $\mu_N$ and $\mu_M$, the average nascent and mature copy numbers (Table S1). If either of these averages exceeded 1,000, we discarded and regenerated the rate vector; we adopted this rejection procedure to ensure the training distributions matched the low-copy number regime seen in transcriptomic data. Next, we evaluated the solution in Equation 18 up to the bounds $\mu_z + 20\sigma_z$ using adaptive quadrature (implemented in the `scipy` function `integrate.quad_vec` [31]) and the inverse real fast Fourier transform (implemented in the `scipy` function `fft.irfft2` [32]). We denote this high-precision procedure by QV20 and treat it as ground truth. After evaluating $P(n, m; \theta)$ on a large grid, we truncated this ground truth PMF to the more practical size $\mathfrak{s}_z = \mu_z + 4\sigma_z$ and stored it to disk. To generate conditional training data, we extracted the entries corresponding to each $n$ and divided them by $P(n; \theta)$ (Equation 7, implemented in the `scipy` function `stats.nbinom.pmf` [31] with $r = \beta^{-1}$ and $p = (1 + b)^{-1}$ [33]). The resulting conditional distributions were stored along with their values of $\theta$ and $n$.

When testing approximations against lower-order generating function solutions, we also calculated generating function solutions using adaptive quadrature on grids of 4 and 10 standard deviations above nascent and mature means, or within the bounds $\mu_z + 4\sigma_z$ and $\mu_z + 10\sigma_z$ in each dimension $z \in \{N, M\}$. We refer to these solutions as QV4 and QV10 in the main text and Figure 2. We also calculated a generating function solution uses 60th-order fixed Gaussian quadrature on a grid of 4 standard deviations above the means (referred to as FQ in the main text and Figure 2).

| Symbol | Definition |
|---|---|
| $K$ | Number of approximating kernels |
| $k$ | Approximating kernel index |
| $\mathbf{w} := w_1, ..., w_K$ | Weights of approximating kernels |
| $\Theta_k$ | Parameters of approximating kernel |
| $h$ | Scaling Factor |
| $\mathcal{F}$ | Generic multivariate function implemented through a neural network |
| $P_{\text{ker}}$ | Univariate PMF of approximating kernel |
| $Q$ | Approximating mixture |
| $\mathfrak{s}_N$ | Nascent state space size or grid dimension |
| $\mathfrak{s}_M$ | Mature state space size or grid dimension |
| $\mathfrak{s}_z$ | State space size in a generic dimension |
| $\mathfrak{s} = \mathfrak{s}_N \times \mathfrak{s}_M$ | Total state space size |
| $P_{\text{NB}}(x; \mu, \sigma)$ | Negative binomial PMF |
| $P_{\text{Poiss}}(x; \lambda)$ | Poisson PMF |

Table S2: Approximation variables.

#### S1.7 Kernel weight regression training and model choice

The default architecture for the neural network $\mathcal{F}$ used to predict $\mathbf{w}$ and $h$ is a multilayer perceptron (MLP) that takes as input rate vector $\tilde{\theta} \in \mathbb{R}^7$ and outputs weight vector $\mathbf{w} \in \mathbb{R}^{10}$, corresponding to 10 kernel functions, and scaling factor $h$. The input $\tilde{\theta}$ is passed through two, fully connected layers of 256 nodes, each with sigmoid activation functions, to output $\mathbf{w}$, which is normalized by the softmax activation function to ensure weights $\mathbf{w}$ sum to unity. To produce $h$, outputs from the first 256 node layer are also passed to a single node with the sigmoid activation function to produce a number in the range [0,1], which is then re-scaled to the range [1,6]. Model architecture is illustrated in Figure S1 and network parameters listed in Table S3.

Weights $\mathbf{w}$, scaling factor $h$, and kernel negative binomial distributions are combined to predict an approximate conditional probability $Q_{\text{KWR}}(m|n; \theta)$ as described in Section S1.3. The network is trained to minimize truncated Kullback-Leibler divergence (Section S1.4) between the approximation $Q_{\text{KWR}}(m|n; \theta)$ and the QV20 conditional ground truth (Section S1.6). We implemented the MLP in PyTorch and used the Adam optimizer to update node weights with a learning rate of 0.001 [34].

Under these conditions, the model successfully decreased the KL divergence between ground truth and approximated conditional distributions for training and validation sets during the training process (Figure S3**A**). The trained model also performed well on a testing set of 756 rate vectors (104,954 conditional examples), with 1% and 99% quantiles of testing KL divergence values of $3.06 \times 10^{-6}$ and 0.0268, respectively. This network required 10.8 hours to train for 35 epochs on a CPU.

To choose our final architecture, we tested the effects of changing model and training configurations on timing and KL divergence minimization, as shown in Figure S3**B-E**. Wherever not indicated, we used 10 kernel functions and two hidden layers of 256 nodes as defaults with 1,536 training rate vectors (211,515 conditional examples), 512 validation rate vectors (72,906 conditional examples), and 768 testing rate vectors (109,685 conditional examples). Models in Figure S3**C** and **D** were trained for 30 epochs, and models in Figure S3e were trained for 35 epochs. Models in Figure S3**B** were trained for 1 epoch with the indicated number of training conditional examples and 1,000 validation conditional examples. Training time scaled linearly with the training set size and increased with the number of kernel functions (Figure S3**B**).

Increasing the number of nodes in each layer allows the network to learn more complex representations: we found that increasing the number of nodes in each hidden layer decreased the testing KL divergences, with diminishing returns at 64 nodes (Figure S3**C**). The number of kernel functions also governs the diversity of approximable distributions: adding more kernels did not produce a substantial effect beyond $K = 10$, which we implement in our final architecture (Figure S3**D**). As we increased the training set size, we found performance improved only marginally (Figure S3**E**). This suggests the function being approximated by $\mathcal{F}$ is relatively simple: the network can be trained on a relatively small number of examples, yet effectively extrapolate to examples outside of the training set.

##### S1.8 Parameter scaled kernel weight regression

Parameter scaled kernel weight regression models were tested and compared for optimal reconstruction of ground truth distributions, as described in Section 2.3. Models $\mathcal{F}_{\text{psKWR}}$ take in vectors $\tilde{\theta}_{\text{psKWR}} \in \mathbb{R}^{4+2K}$ where $K$ is the number of kernel functions, as described in Section S1.5.

Inputs $\tilde{\theta}_{\text{psKWR}}$ were passed through several fully connected layers to produced weights $\mathbf{w} \in \mathbb{R}^K$, initial mean scaling factors $\mathbf{s}_\mu \in \mathbb{R}^K$, and initial standard deviation scaling factors $\mathbf{s}_\sigma \in \mathbb{R}^K$. Weights $\mathbf{w}$ had the softmax function applied to sum to unity, and scaling vectors $\mathbf{s}_\mu$ and $\mathbf{s}_\sigma$ had sigmoid activation functions applied to sum to 0 to 1. Then, scaling factors were multiplied by model parameter $C$ such that scaling vectors modified approximated means by at most a factor of $C$. $C$ was initialized at 2 and updated during training.

Models were trained to minimize Hellinger distance between joint microstate probabilities (Section S1.4).

For each of 3, 5 and 10 kernel functions, various model architectures were tested. With the number of nodes held constant at 256 per layer, models of 1, 2, 3, 4, and 5 hidden layers with ReLU activation functions, the output of the which were passed separately to a layer for kernel weights with a softmax activation function, a scaling factors for kernel means with a sigmoid activation function, and a layer for scaling factors for kernel standard deviations with a sigmoid activation function were trained. Models were trained with 64, 128, 256, and 512 nodes per layer with constant 2 hidden layers with ReLU activation functions. All models were trained for 100 epochs 1,280 rate vectors $\theta$ (corresponding to 211,515 conditional vectors $\tilde{\theta}_{\text{psKWR}}$) and 512 validation rate vectors $\theta$ (corresponding to 72,906 conditional vectors $\tilde{\theta}_{\text{psKWR}}$). Testing results for 49,704 testing rate vectors (with measures of KL divergence, Hellinger distance, and Mean Squared Error against QV20 ground truth solutions) motivate final psKWR model architecture. Models were implemented in Pytorch and optimized via the Adam optimizer with a learning rate of 0.001 [34].

The final network architecture for psKWR approximations comprises 2 hidden layers before the output layers of 512 nodes per layer to produce weights and scaling factors for 5 kernel functions.

It is diagrammed in Figure S2 with layers and parameters summarized in Table S4.

#### S2 Direct Regression (DR)

In Figure S5, we compare the performance of the proposed procedure to a method more typical of machine learning: using a neural network to directly compute the joint probability given a rate vector and the number of nascent and mature RNA molecules: $\mathcal{G} : \theta_D \mapsto Q(n, m; \theta)$. Here, a direct example $\theta_D = \{\log_{10} b, \log_{10} \beta, \log_{10} \gamma, n, m\}$. We refer to this method as direct regression (DR). Here, the neural network directly approximates the generating function solution, bypassing the need for distributional approximations and kernel functions. Dense, fully connected MLPs ranging from 128 to 3000 nodes and 2 to 5 layers with ReLu activation functions between each layer were used for direct regression. Models were trained on 512 or 1,024 training rate vectors $\theta = \{b, \beta, \gamma\}$ corresponding to 14,027,057 and 24,481,592 direct examples respectively, with 700,000 validation direct examples. Models were implemented in PyTorch using the Adam optimizer with a learning rate of 0.001 and weight decay of 0.0001 [34] and trained for 20 epochs to reduce the mean squared error between the predicted probabilities $Q_{\text{DR}}(n, m; \theta)$ and $P(n, m; \theta)$. After training architectures with different number of layers and nodes, we tested performance by predicting probabilities over state spaces and comparing direct regression approximations to QV20 distributions (ground truth, described in Section S1.6). Figure S5 shows Hellinger distances between direct approximation and ground truth (QV20) for 756 testing rate vectors. The model used for comparison in Figure 2 was chosen based on results in Figure S5 and comprises 3 layers of 256 nodes. It was trained on 24,481,592 examples (1,024 rate vectors) for 20 epochs. Parameters for the final model are listed in Table S5.

#### S3 Moment-matched lognormal/NB approximation procedure (MMNB)

As discussed in Section S1.5, we use a moment-matched lognormal distribution to coarsely approximate the location and dispersion of each conditional distribution $P(m|n)$. We can go one step further and dispense with the neural network altogether, plugging these parameters back into a conditional negative binomial distribution. This gives us the following PMF:

$$\begin{aligned} Q_{MM}(n, m) &= P(n)Q(m|n), \\ Q(m|n) &= P_{NB}(\mu_{2,n+1}, \sigma_{2,n+1}), \\ \mu_{2,n+1} &= \exp\left(\mu_{yl,n+1} + \sigma_{yl,n+1}/2\right), \\ \sigma_{2,n+1} &= \mu_{2,n+1}\sqrt{\exp\left(\sigma^2_{yl,n+1}\right) - 1}, \end{aligned} \tag{37}$$

where the two final equations are standard properties of the lognormal distribution, with their arguments computed through Equation 16.

#### S4 Supplementary Figures and Tables

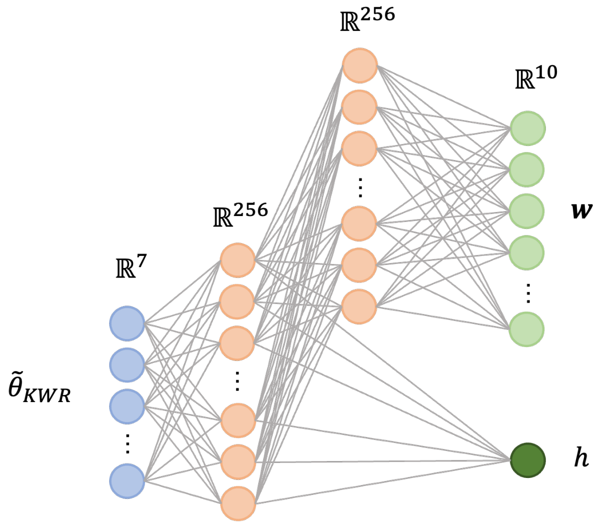


Figure S1: Architecture of the final model for KWR. It includes an input layer, two fully connected layers of 256 nodes and output layer of 10 nodes that produces weights for kernel functions. Another layer produces the scaling factor $h$ with outputs from first layer of 256 nodes.

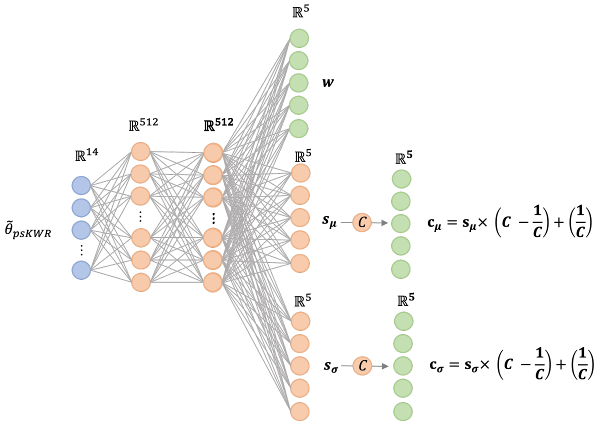


Figure S2: Architecture of the final model for psKWR. It includes an input layer, two fully connected layers of 512 nodes with ReLu activation functions whose outputs are input to one layer with a softmax activation function that produces weights for kernel functions, one output layer of 512 nodes that produces scaling factors for 5 kernel means, and another output layer that produces scaling factors for 5 kernel standard deviations. Scaling factors are adjusted by (the same, though illustrated twice) network parameter C to produce final scaling factors (see Section S1.8).

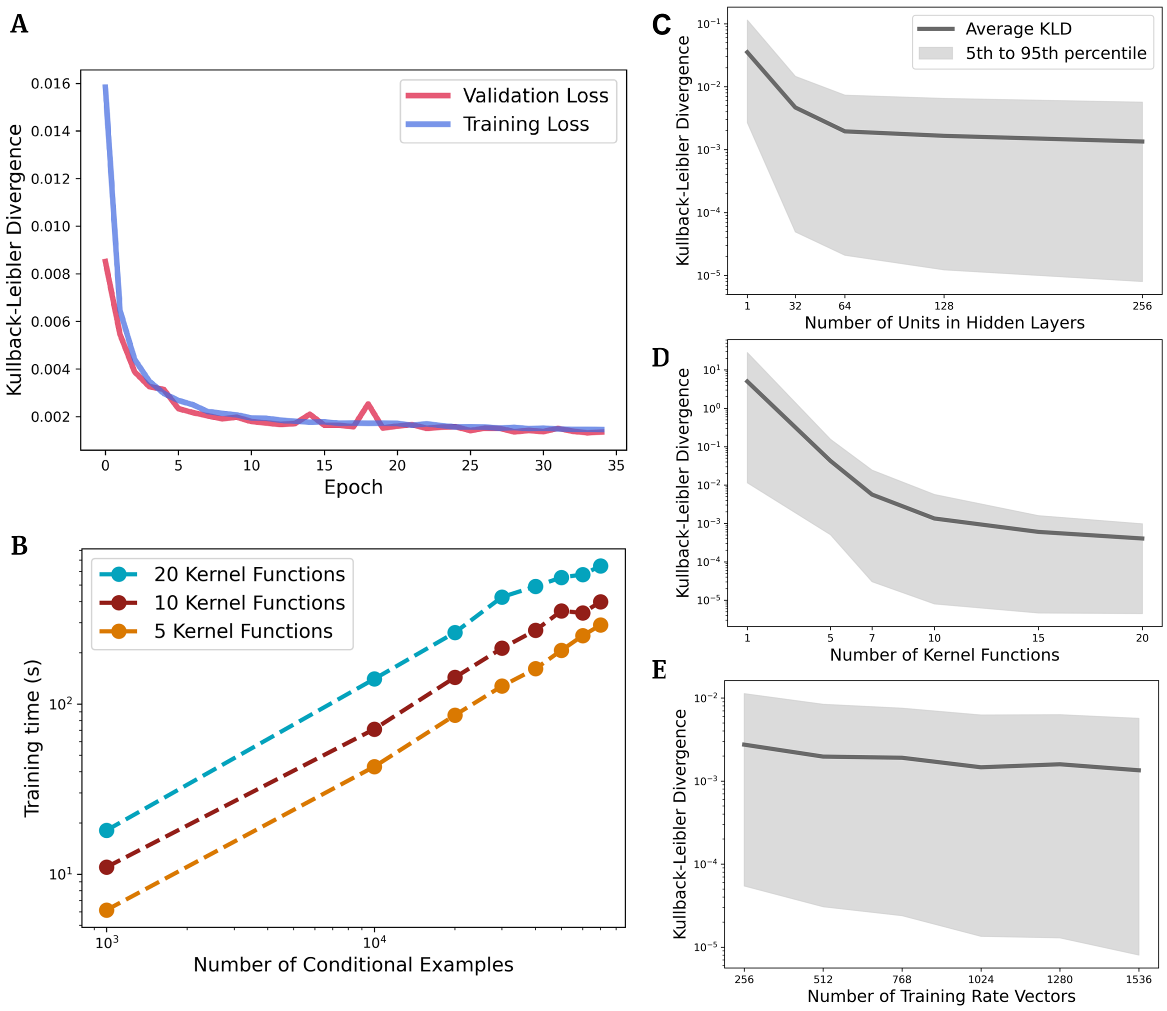


Figure S3: The training, validation, and testing performance of the KWR method. **A.** The average Kullback-Leibler divergence (KL divergence) between true and KWR approximated conditional distributions for training and validation examples decreases to zero as the model is trained. **B.** The training time increases linearly with the size of the training set and the number of kernel functions used. **C.** KL divergences for testing examples decrease with increasing the number of nodes in hidden layer, **D.** increasing the number number of kernel functions, and **E.** increasing the number of training rate vectors.

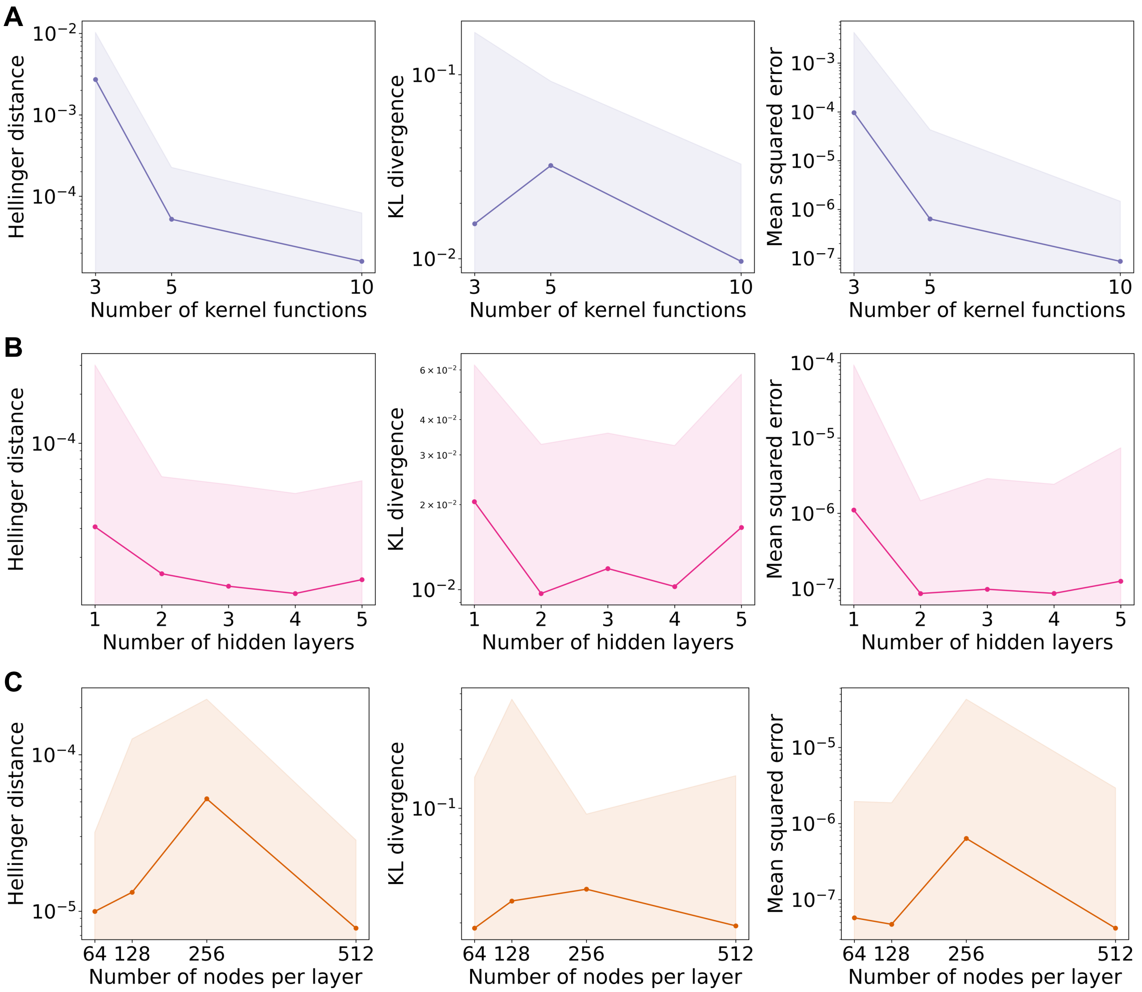


Figure S4: Comparison of testing results for different psKWR model architectures. Hellinger distance, truncated Kullback-Leibler divergence (KL divergence) (see Section S1.4), and Mean Squared Error between psKWR approximated distributions and QV20 ground truth solutions for 49,704 testing conditional examples ($\tilde{\theta}_{\text{psKWR}}$). Distances as a function of **A.** varying number of kernel basis functions, **B.** varying numbers of hidden layers, and **C.** varying numbers of nodes per layer.

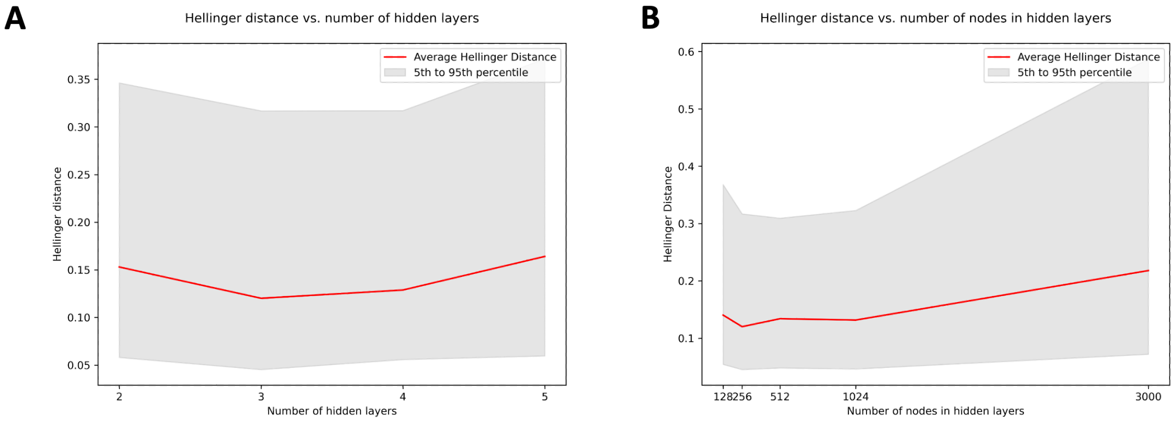


Figure S5: Performance and comparison of direct regression architectures: Hellinger distance between generating function based solutions (QV20) and direct regression approximations for 768 testing rate vectors. Approximations did not improve with increasing **A**. number of layers (with a fixed 256 nodes per layer) or **B**. number of nodes in hidden layers (with a fixed 3 layers). Best accuracy (lowest Hellinger distance) was for an architecture of 3 fully connected layers of 256 nodes each.

| Name | Number of Parameters | Activation Function | Input |
|---|---|---|---|
| L1 | 2,048 | sigmoid | $\tilde{\theta}_{\mathrm{KWR}}$ |
| L2 | 65,792 | sigmoid | L1 |
| L3: Kernel weights | 2,570 | softmax | L2 |
| L4: Scalar for standard deviation | 257 | sigmoid | L1 |

Table S3: Parameters of the final network used for kernel weight regression (KWR). Total trainable parameters: 70,667. Input is $\tilde{\theta}_{\mathrm{KWR}}$ described in Section S1.7.

| Name | Number of Parameters | Activation Function | Input |
|---|---|---|---|
| L1 | 7,168 | ReLU | $\tilde{\theta}_{\mathrm{psKWR}}$ |
| L2 | 262,144 | ReLU | L1 |
| L3: Kernel weights | 3,702 | softmax | L2 |
| L4: Scaling vector for kernel means | 3,702 | sigmoid | L2 |
| L5: Scaling vector for kernel standard deviations | 3,702 | sigmoid | L2 |

Table S4: Parameters of the final network used for parameter scaled kernel weight regression (psKWR). Total trainable parameters: 280,418. Input is $\tilde{\theta}_{\mathrm{psKWR}}$ described in Section 2.3.

| Name | Number of Parameters | Activation Function | Input |
|---|---|---|---|
| L1 | 1,536 | ReLu | $\theta_D$ |
| L2 | 65,792 | ReLu | L1 |
| L3 | 65,792 | ReLu | L2 |
| L4 | 257 | sigmoid | L3 |

Table S5: Parameters of the final network used for direct regression (DR). Total trainable parameters: 133,377. Input is $\theta_D$ described in Section S2.
